# Strengths and limitations of using participatory science data to characterize a wildlife mass mortality event

**DOI:** 10.1101/2024.05.02.592273

**Authors:** Liam U. Taylor, Tatsiana Barychka, Seabird McKeon, Natasha Bartolotta, Stephanie Avery-Gomm

## Abstract

Large participatory science (i.e., “community science” or “citizen science”) platforms are increasingly used at every level of ecological and conservation research, including disease monitoring. Here, we used a comprehensive, ground-truthed mortality dataset to judge how well participatory science data from iNaturalist represented the magnitude, taxonomic, temporal, and spatial patterns of waterbird mortality associated with a mass mortality event following the incursion of Highly Pathogenic Avian Influenza in eastern Canada in 2022. The iNaturalist dataset was effective at identifying species with high mortality (especially Northern Gannets, *Morus bassanus*), along with the time period and spatial regions with high concentrations of avian deaths. However, iNaturalist data severely underestimated the magnitude, overestimated the taxonomic breadth, and poorly represented the full geographic scope of disease-related deaths. Our results suggest iNaturalist can be used to identify the species, timing, and location of relatively high mortality in situations where no other information is available, and to supplement conventional sources of data. However, iNaturalist alone can neither quantify the magnitude nor pinpoint the mechanisms of mortality and therefore is not a viable substitute for comprehensive mortality assessments.

## INTRODUCTION

Participatory science (also known as “community science” or “citizen science”) is broadly defined as scientific research or knowledge produced with public participation. Participatory science now plays a key role in many ecological research programs, from environmental quality monitoring (Conrad and Hilchey 2011), to species distribution modeling (Feldman et al. 2021), to research on wildlife disease and mortality (Lawson et al. 2015). For example, Altizer et al. (2004) leveraged decades of volunteer observations to analyze finch conjunctivitis in eastern North America, while Szentivanyi and Vincze (2022) used photographs from iNaturalist and other public repositories to identify the spread of toad myiasis in Europe. Online, opportunistic participatory science platforms such as iNaturalist are increasingly used to investigate mortality or wildlife-related disease across a wide range of organisms (Yang et al. 2021, Cull 2022, Bartolotta et al. 2023, Rhodes et al. 2023).

The avian panzootic of Highly Pathogenic Avian Influenza (HPAI) virus H5N1 2.3.4.4b is estimated to have already killed hundreds of thousands, if not millions, of wild birds (Klaassen and Wille 2023). Recent HPAI outbreaks raise new concerns about not only wild bird conservation (UNEP 2023) but also deadly transmission events into terrestrial and marine mammals (e.g., Leguia et al. 2023, Elsmo et al. 2023). It is thus critical to monitor the spread and impact of this virus, which is now most apparent in wild birds through mass die-off events (e.g., Rijks et al. 2022). In facing the global spread of HPAI, however, practitioners in some regions may have a limited capacity to collect disease-related data and monitor mortality.

The objective of this study was to evaluate how well an accessible, participatory science dataset from iNaturalist—in the absence of additional data—represented mortality during a regional wave of HPAI in wild bird populations in eastern Canada. Using a comprehensive dataset of disease-related avian mortality (Avery-Gomm et al. 2024 [preprint]), we ask how effectively an iNaturalist dataset represented the (1) magnitude, (2) taxonomic distribution, (3) temporal patterns, and (4) spatial distribution of wild bird mortality associated with the incursion of HPAI.

## METHODS

### Datasets

We compared two datasets: (1) a comprehensive, heavily-curated, dataset of HPAI-linked mortality, and (2) an iNaturalist dataset, representing the kind of information that would be available without curation, supplementation, or ground-truthing. Both datasets cover the same study area and time period. The study area spans the Canadian provinces of Quebec, Newfoundland and Labrador, New Brunswick, Nova Scotia, and Prince Edward Island, along with the French territory of Saint Pierre et Miquelon. The time period runs from 1 April to 30 September, 2022.

We used the comprehensive HPAI-linked mortality dataset from Avery-Gomm et al. (2024 [preprint]). This dataset is compiled from federal, provincial, Indigenous, and municipal government databases, the Canadian Wildlife Health Cooperative and other NGOs, academic researchers, and a small proportion of records from two participatory science platforms (iNaturalist and eBird). The comprehensive dataset is pre-filtered to exclude double-counted records (“Scenario B”) and exclude records from taxa that were not presumed to be positive for HPAI based on regional testing results (e.g., Leach’s Storm-Petrels, *Hydrobates leucorhous*, which were found dead but never HPAI-positive; Giacinti et al. 2023 [preprint]). Avery-Gomm et al. (2024 [preprint]) are clear this published dataset has its own limitations—representing only a subset of total mortality in the region—but in this paper, we treat it as complete.

For the iNaturalist dataset, we used the iNaturalist project titled “HPAI|Dead birds in Eastern Canada during the 2022 HPAI outbreak.” Data was extracted on 6 January 2023. This project contains all records of photographed birds annotated as “dead” by iNaturalist users, including both “Research Grade” and “Needs ID” records. We excluded seven “Needs ID” records that featured only photos of individual bleached bones. For this study, we used iNaturalist because it is an opportunistic platform that encourages users to upload individual wildlife observations, including carcasses. Although a different platform, eBird, provides the majority of participatory science data about birds, eBird is built around species checklists that are useful for studies of population range or abundance as opposed to individual mortality (Sullivan et al. 2014, Neate-Clegg et al. 2020).

We filtered both datasets to include only waterbirds spanning six orders— Anseriformes, Charadriiformes, Suliformes, Procellariiformes, Pelecaniformes, and Gaviiformes—following the Clements v2023 taxonomy (Clements et al. 2023). Both datasets originally featured taxonomically ambiguous records (e.g., “*Larus*”, “Alcidae”, “Charadriiformes”). We retained records if they were identifiable to at least taxonomic family.

Note that curated reports from iNaturalist and eBird are included in the comprehensive dataset but represent only a small proportion of records (473/40,109 mortalities, 279/3,728 records). Avery-Gomm et al. (2024 [preprint]) describe the curation process for the iNaturalist data in the comprehensive dataset. The iNaturalist dataset used in this study is not curated, as our goal is to evaluate the effectiveness of participatory science data precisely in the absence of outside knowledge.

### Magnitude and taxonomic comparisons

First, we compared the magnitude and taxonomic distribution of avian mortality as represented in the comprehensive and iNaturalist datasets. Specifically, we asked three questions: (1) Does the iNaturalist dataset represent the absolute magnitude of avian mortality? (2) Does the iNaturalist dataset capture the relative distribution of mortality across different taxonomic groups of birds? (3) Does the iNaturalist dataset provide a precise taxonomic representation of HPAI-specific mortality, as opposed to avian mortality in general?

### Temporal comparisons

Next, we compared the timing of mortality records between datasets. Specifically, we asked how precisely and accurately the iNaturalist dataset identified the core period and peak of mortality as judged by the comprehensive mortality dataset. For each dataset, we defined: (1) core period, as the shortest interval containing the 51% (majority) of reported mortality; and (2) peak, as the week of maximum reported mortality. We binned records by week for these comparisons.

### Spatial comparisons

Finally, we compared the spatial distribution of mortality reported in each dataset. Specifically, we asked how accurately iNaturalist data captured the regional extent and local concentrations of avian deaths. In analogy to methods from movement ecology such as home range analysis, we define spatial distribution as the 95% isopleth of mortality locations (i.e., the regions containing 95% of mortality locations as estimated from underlying kernel density). We estimated kernel density with the *kernelUD* function (grid size 1000, extent 1, shared grid) and 95% isopleths with the *getverticeshr* function from R package *adehabitatHR* (Calenge 2023). Geolocation precision differed between and within datasets, so we rounded all latitude and longitude values to two decimal points.

Each location record in the iNaturalist dataset represents one dead bird. In contrast, records from the comprehensive dataset represent 1–5,119 deaths at a given location. We thus quantified two separate spatial distributions from the comprehensive dataset. The comprehensive “occurrence” distribution ignores the number of deaths and treats each record in the comprehensive dataset as a single point. The comprehensive “concentration” distribution duplicates records in the comprehensive dataset so that each point represents an individual dead bird (e.g., a record of ten dead birds in the comprehensive dataset results in ten records of one bird). Whereas the “occurrence” distribution helps estimate the full geographic range of HPAI-linked mortality, the “concentration” distribution emphasizes areas with high numbers of avian deaths. Using the *kerneloverlap* function (method=“HR”) from *adehabitatHR*, we report the proportion of comprehensive “occurrence” and “concentration” spatial distributions in the comprehensive dataset that are covered by the iNaturalist spatial distribution.

Quantitative spatial comparisons were sensitive to the smoothing parameter, *h*, used for estimating kernel densities: higher *h* values lumped all locations into a single, vast contour spanning the region (see Figure S1 for results across a range of *h* values, 0.1–0.9). Our main analysis used a relatively small value, *h* = 0.3, because this value smoothed over adjacent records in the comprehensive mortality occurrence distribution while still restricting occurrence areas to plausible coastal habitats for waterbirds. Data were wrangled and visualized using the *sf* and *tidyverse* packages in R version 4.3.2 (Pebesma 2018, Wickham et al. 2019, R Core Team 2023).

## RESULTS

### Magnitude and taxonomic comparisons

The iNaturalist dataset severely underestimated the overall scale of waterbird mortality during the course of HPAI outbreaks in eastern Canada during spring and summer 2022. The comprehensive dataset reported 40,109 dead waterbirds linked to HPAI across 3,728 records. The iNaturalist dataset contained only 283 records (90 observers, 1–65 records per observer), each of which was assumed to represent a single dead bird.

The iNaturalist dataset was generally accurate in representing the relative taxonomic distribution of avian mortality (Fig. 1). The most commonly reported species was Northern Gannet (*Morus bassanus*; iNaturalist: N = 143/283 reported mortalities; comprehensive: N = 25,669/40,109 reported mortalities; Fig 1A, Appendix S1: Table S1). The second most common species was the Common Murre (Alcidae: *Uria aalge*: iNaturalist: N = 40/283 reported mortalities; comprehensive: N = 8,133/40,109 reported mortalities). Both datasets were also generally consistent in moderate mortality across four families—gulls/terns (Laridae), auks (Alcidae), waterfowl (Anatidae), and cormorants (Phalacrocoracidae)— although Laridae was relatively underreported in the iNaturalist dataset (Fig. 1B). Both datasets featured additional reports from shearwaters (Procellariidae), herons (Ardeidae), and sandpipers/woodcocks (Scolopacidae; Fig. 1B, Appendix S1: Table S1).

**Figure 1.**
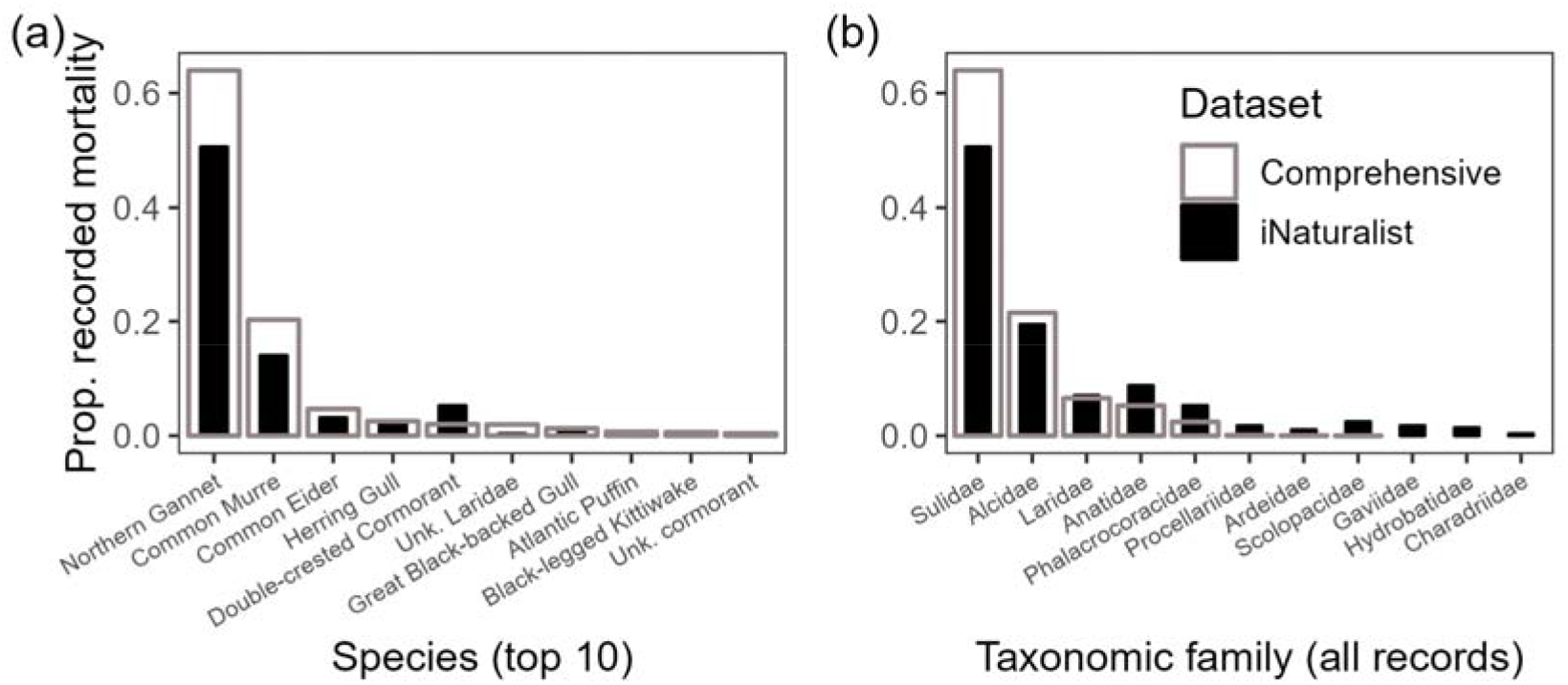
Taxonomic comparison of avian deaths between iNaturalist data and a comprehensive dataset of HPAI-linked mortality in waterbirds, eastern Canada 2022. Gray bars show the proportion of deaths among different waterbird taxa within the comprehensive dataset. Black bars show the proportion of deaths within the smaller iNaturalist dataset. (a) Organized by species, showing only the 10 most commonly reported taxa from the comprehensive dataset; (b) Organized by taxonomic family, showing all records. See Table S1 for original taxonomic labels.

There were both species- and family-level errors in the taxonomic coverage of the iNaturalist dataset. At the species level, the iNaturalist dataset missed taxa. There were zero records of dead Atlantic Puffins (Alcidae: *Fratercula arctica*) and Great Shearwaters (Procellariidae: *Ardenna gravis*) in the iNaturalist dataset, despite these species having 282 and 34 mortalities in the comprehensive dataset, respectively (Appendix S1: Table S1). At the family level, the iNaturalist dataset overestimated the scope of HPAI-related mortality by including records of loons (Gaviidae), storm-petrels (Hydrobatidae), and plovers (Charadriidae; (Fig. 1B). Mortality in these taxa was excluded from the comprehensive dataset because no individuals in these taxonomic groups tested positive for HPAI in the study area during the study period (Giacinti et al. 2023 [preprint]); and therefore Avery-Gomm et al. (2024 [preprint]) did not attribute mortalities to HPAI.

### Temporal comparisons

The iNaturalist dataset accurately identified the core period of mortality. The core period, defined as the shortest interval containing a majority (51%) of recorded mortality, spanned four weeks in the iNaturalist dataset (28 May to 25 June), falling directly within the core period of the comprehensive dataset (six weeks; 28 May to 9 July; Fig. 2A).

**Figure 2.**
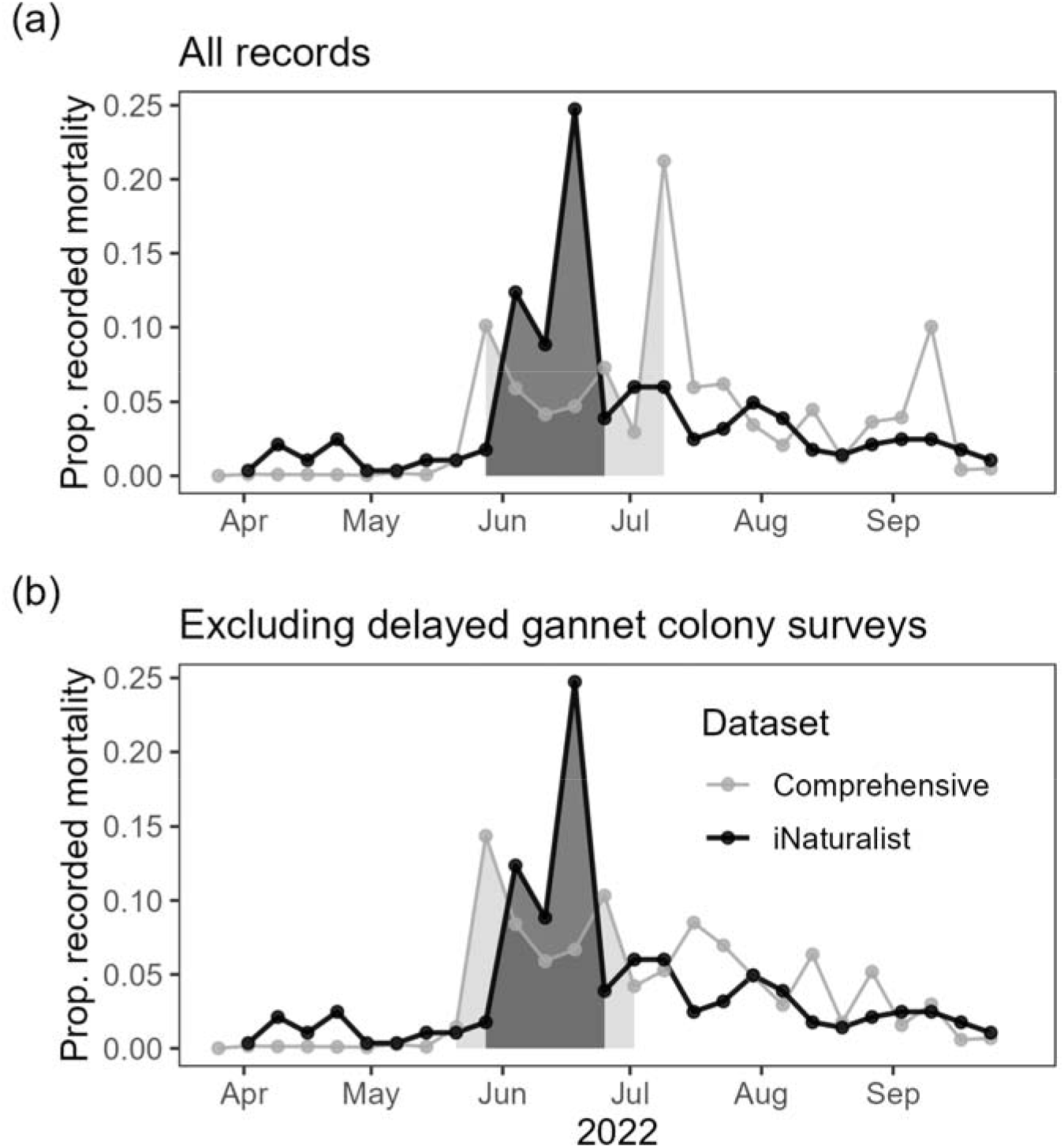
Temporal comparison of avian deaths between iNaturalist data and a comprehensive dataset of HPAI-linked mortality in waterbirds, eastern Canada 2022. Records are binned by week. Shaded regions mark the shortest interval containing 51% of record mortalities. (a) Weekly proportion of recorded mortality. (b) Weekly proportions after excluding gannet colony surveys, which offered delayed snapshots of mortality, in the comprehensive dataset.

The iNaturalist dataset initially failed to identify weeks of peak mortality in the comprehensive dataset (Fig. 2A). The week of maximum mortality from iNaturalist was 18 June (70 mortalities; 8 observers, 1–53 records per observer). In contrast, the week of maximum mortality from the comprehensive dataset was 9 July (8,518 deaths), with a second peak the week of 10 September (4,028 deaths). However, Avery-Gomm et al. (2024 [preprint]) note these peaks in the comprehensive dataset are influenced by a reporting lag related to the timing of Northern Gannet colony surveys by Environment and Climate Change Canada (ECCC) staff. When we removed ECCC gannet colony surveys from the comprehensive dataset, the week of peak mortality shifted to 28 May. The peak reported by iNaturalist then falls three weeks behind the comprehensive peak (Fig. 2B).

### Spatial comparisons

The spatial distribution of iNaturalist mortality records covered most regions of HPAI-related mortality as represented in the comprehensive dataset (Fig. 3). The iNaturalist distribution covered 70% of the mortality concentration distribution, with the remaining high-concentration areas falling just outside of iNaturalist boundaries (Fig. 3). The iNaturalist dataset covered a somewhat smaller proportion (61%) of the mortality occurrence distribution, with several occurrence areas in the Gulf of Saint Lawrence, the coast of Quebec, and the coast of Labrador not represented by locations in the iNaturalist dataset (Fig. 3). Overlap statistics varied widely depending on the *h* parameter used to estimate spatial distributions. The iNaturalist spatial distribution only covered ∼30–40% of the comprehensive distributions at *h* = 0.1 (low smoothing), whereas coverage reached ∼80% at *h* = 0.9 (high smoothing; Appendix S1: Fig. S1). However, overall patterns were consistent across all sampled *h* values: the iNaturalist spatial distribution covered, or fell adjacent to, the bulk of the comprehensive concentration distribution while failing to cover some northern portions of the comprehensive occurrence distribution.

**Figure 3.**
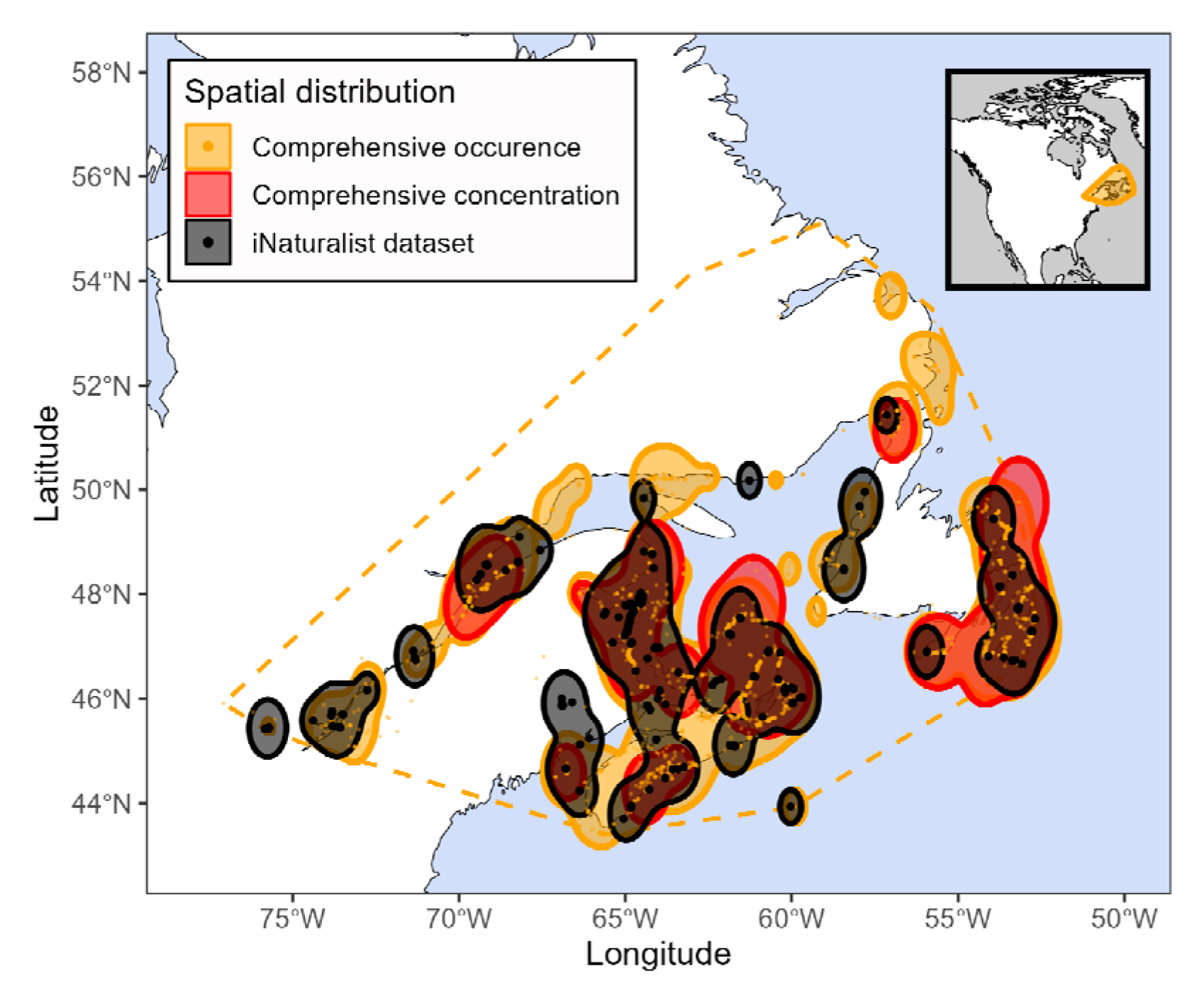
Spatial comparison of avian deaths between iNaturalist data and a comprehensive dataset of HPAI-linked mortality in waterbirds, eastern Canada 2022. Spatial distribution is defined as the 95% isopleth of mortality locations (kernel smoothing *h* = 0.3). Black contours show spatial distribution from the iNaturalist dataset. Comprehensive occurrence distribution (orange) treats mortality records from the comprehensive dataset as individual points, regardless of reported mortality, whereas the comprehensive concentration distribution (red) duplicates mortalities based on the number of dead birds reported at each location. Orange dashed line indicates the full extent (convex hull) of mortality reports from the comprehensive dataset.

## DISCUSSION

Our comparison of a participatory science dataset to a comprehensive dataset of HPAI-related mortality among waterbirds in eastern Canada, 2022, demonstrated both general strengths and fundamental limitations in using iNaturalist data to draw conclusions about a mass mortality event. In terms of magnitude, the iNaturalist dataset included only 283 records of dead waterbirds in the region, less than 1% of avian deaths estimated from the 2022 outbreak (Appendix 1: Table S1; Avery-Gomm et al. 2024 [preprint]). In terms of taxonomic coverage, the iNaturalist dataset accurately identified species with the largest recorded numbers of HPAI-related mortality (Northern Gannets and Common Murres; Fig. 1A) and successfully reported mortality across eight waterbird families known to have been affected by HPAI (Fig. 1B). Yet iNaturalist records missed some species that were documented in the comprehensive mortality dataset, such as Atlantic Puffins (Appendix S1: Table S1). More critically—if the intent is to monitor HPAI-linked mortality—iNaturalist included mortality records from entire taxonomic families such as loons (Gaviidae) and storm-petrels (Hydrobatidae; Fig. 1B), though these deaths were likely not associated with HPAI (Giacinti et al., 2023 [preprint]).

In terms of temporal patterns, the iNaturalist dataset precisely identified the core period of mortality associated with the incursion of HPAI in May–July 2022 (Fig. 2A). Indeed, the week of peak mortality differed by only three weeks between datasets once delayed seabird colony survey reports were excluded from the comprehensive dataset (Fig. 2B). In terms of spatial distribution, iNaturalist were congruent with the comprehensive dataset, either covering or bordering all high-concentration mortality areas (Fig. 3). However, there were zero iNaturalist records to represent some outbreak occurrence regions in parts of Labrador and the Gulf of St. Lawrence (Fig. 3).

Although we are emphasizing the limitations of using this simplest form of iNaturalist data, there are several ways to improve the application of these data for wildlife mortality monitoring. The participatory science dataset here is drawn from a single source (iNaturalist) with no curation or adjustments beyond basic taxonomic assignments. For example, Avery-Gomm et al. (2024 [preprint]) describe two methods to uncover unmarked records of dead birds from iNaturalist (i.e., screen observations by known volunteer observers conducting beach surveys and extract all records of species with the highest reported mortality to manually check annotations). Their curation efforts also included excluding mortality records for species where deaths could not be attributed to HPAI. Such curation improved the contributions of participatory science data to the comprehensive dataset, but only by leveraging information from outside of participatory science itself.

Further, participatory science records can be heavily biased toward urban, accessible, or historically wealthy areas (Tiago et al. 2017, Bowler et al. 2022, Ellis-Soto et al. 2023). Accordingly, the iNaturalist dataset in this study lacked mortality records from less-accessible areas along the northern coast of the Gulf of Saint Lawrence and in Labrador (Fig. 3). Application-specific models can help correct participatory science records for biological and observer effort patterns (e.g., Callaghan et al. 2022). However, the long-term or ground-truth information needed for accurate model corrections may be exactly what is unavailable to a practitioner forced to rely on sparse participatory science data for wildlife monitoring in the first place.

Mass mortality events associated with the HPAI virus are on the rise, and assessing mortality is an important but complex objective of monitoring outbreaks (Ramey et al. 2022, Klaassen and Wille 2023). Previous work has envisioned a combination of monitoring pipelines that together enable comprehensive disease surveillance akin to a One Health approach (e.g., Morner et al. 2002, Ramey et al. 2022, Cardoso et al. 2022, Sharan et al. 2023). Each monitoring pipeline has its strengths and weaknesses. We cannot overstate the importance of proactive disease testing programs, which identify the mechanisms of mortality and may also operate as early-warning systems for emerging outbreaks (Kelly et al. 2021). Yet laboratory testing alone can only offer data on a subset of affected wildlife (Klaassen and Wille 2023). Government agencies can organize thorough population surveys or visits to otherwise inaccessible areas, but these are often slow, expensive, and sporadic (e.g., the individual colony reports that appeared as mortality peaks in Fig. 2A). Online participatory science databases such as iNaturalist are freely-available, updated continuously, and cover global scales, but may provide sparse or biased mortality records (Theobald et al. 2015, Lawson et al. 2015). Participatory science data will be most useful as part of a broader, integrated effort to assess mortality that includes all of these pipelines.

In conclusion, our study suggests that iNaturalist data alone can help characterize mass mortality events—whatever the cause—but with major limitations. While relying on iNaturalist data alone would severely underestimate the absolute magnitude of mortality and under-represent the full geographic extent of a mortality event, it may be useful for identifying the main species, time period, and geographic regions of highest mortality. When mass mortality events are associated with HPAI, iNaturalist data cannot directly identify which mortalities can be attributed to HPAI, thus potentially overestimating the taxonomic breadth of host populations. However, these recommendations are based on our single case study. Generalizing these recommendations will require future research that explores how well iNaturalist data can characterize wildlife mortality across multiple geographic regions and multiple causes of death.

## ACKNOWLEDGEMENTS

Funding for this project was provided by the Science and Technology Branch of Environment and Climate Change Canada. We are grateful to Mark Pokras and Caleb Spiegel for feedback on the initial analysis, and Diego Ellis-Soto and Yen-Hua Huang for comments on the manuscript. The idea of using iNaturalist to understand mortality was first discussed among members of the Atlantic Marine Bird Cooperative’s Community Science and Marine Bird Health Working Group.

