## Supplementary Material for "Strengths and limitations of using participatory science data to characterize a wildlife mass mortality event"

**APPENDIX S1**

**Table S1.** Avian mortality in the comprehensive and iNaturalist datasets by original taxonomic label

|  |  |  |  | **Recorded mortality in dataset** | |
| --- | --- | --- | --- | --- | --- |
| **Original label** | **Common name** | **Family** | **Order** | **Comprehensive**  **(total = 40,109)** | **iNaturalist**  **(total = 283)** |
| *Morus bassanus* | Northern Gannet | Sulidae | Suliformes | 25,669 | 143 |
| *Uria aalge* | Common Murre | Alcidae | Charadriiformes | 8,133 | 40 |
| *Somateria mollissima* | Common Eider | Anatidae | Anseriformes | 1,894 | 9 |
| *Larus argentatus* | Herring Gull | Laridae | Charadriiformes | 1,030 | 5 |
| *Nannopterum auritum* | Double-crested Cormorant | Phalacrocoracidae | Suliformes | 813 | 0 |
| Laridae sp. | Unk. Laridae | Laridae | Charadriiformes | 796 | 0 |
| *Larus marinus* | Great Black-backed Gull | Laridae | Charadriiformes | 522 | 3 |
| *Fratercula arctica* | Atlantic Puffin | Alcidae | Charadriiformes | 282 | 0 |
| *Rissa tridactyla* | Black-legged Kittiwake | Laridae | Charadriiformes | 251 | 2 |
| Phalacrocoracidae sp. | Unk. cormorant | Phalacrocoracidae | Suliformes | 168 | 0 |
| *Alca torda* | Razorbill | Alcidae | Charadriiformes | 119 | 5 |
| *Branta canadensis* | Canada Goose | Anatidae | Anseriformes | 107 | 9 |
| *Anas platyrhynchos* | Mallard | Anatidae | Anseriformes | 70 | 4 |
| *Uria lomvia* | Thick-billed Murre | Alcidae | Charadriiformes | 46 | 4 |
| *Alle alle* | Dovekie | Alcidae | Charadriiformes | 46 | 2 |
| *Ardenna gravis* | Great Shearwater | Procellariidae | Procellariiformes | 34 | 0 |
| *Larus delawarensis* | Ring-billed Gull | Laridae | Charadriiformes | 26 | 4 |
| *Anser caerulescens* | Snow Goose | Anatidae | Anseriformes | 24 | 1 |
| *Ardea herodias* | Great Blue Heron | Ardeidae | Pelecaniformes | 15 | 2 |
| *Anas rubripes* | American Black Duck | Anatidae | Anseriformes | 15 | 0 |
| *Sterna paradisaea* | Arctic Tern | Laridae | Charadriiformes | 13 | 0 |
| *Sterna hirundo* | Common Tern | Laridae | Charadriiformes | 10 | 0 |
| *Fulmarus glacialis* | Northern Fulmar | Procellariidae | Procellariiformes | 6 | 2 |
| *Melanitta perspicillata* | Surf Scoter | Anatidae | Anseriformes | 4 | 1 |
| *Larus glaucoides* | Glaucous-winged Gull | Laridae | Charadriiformes | 3 | 0 |
| *Aix sponsa* | Wood Duck | Anatidae | Anseriformes | 2 | 0 |
| *Melanitta americana* | Black Scoter | Anatidae | Anseriformes | 2 | 0 |
| *Mergus merganser* | Common Merganser | Anatidae | Anseriformes | 2 | 0 |
| *Spatula discors* | Blue-winged Teal | Anatidae | Anseriformes | 2 | 0 |
| *Anas acuta* | Northern Pintail | Anatidae | Anseriformes | 1 | 0 |
| *Branta bernicula nigricans* | Brant | Anatidae | Anseriformes | 1 | 0 |
| *Branta hutchinsii* | Cackling Goose | Anatidae | Anseriformes | 1 | 0 |
| *Mergus serrator* | Red-breasted Merganser | Anatidae | Anseriformes | 1 | 0 |
| *Tringa semipalmata* | Willet | Scolopacidae | Charadriiformes | 1 | 0 |
| *Phalacrocorax auritus* | Double-crested Cormorant | Phalacrocoracidae | Suliformes | 0 | 15 |
| *Gavia immer* | Common Loon | Gaviidae | Gaviiformes | 0 | 4 |
| *Hydrobates leucorhous* | Leach's Storm-Petrel | Hydrobatidae | Procellariiformes | 0 | 4 |
| Larinae | Unk. gull | Laridae | Charadriiformes | 0 | 3 |
| *Scolopax minor* | American Woodcock | Scolopacidae | Charadriiformes | 0 | 3 |
| *Ardenna grisea* | Sooty Shearwater | Procellariidae | Procellariiformes | 0 | 2 |
| *Calidris pusilla* | Semipalmated Sandpiper | Scolopacidae | Charadriiformes | 0 | 2 |
| *Uria* | Unk. murre | Alcidae | Charadriiformes | 0 | 2 |
| Alcidae | Unk. auk | Alcidae | Charadriiformes | 0 | 1 |
| *Botaurus lentiginosus* | American Bittern | Ardeidae | Pelecaniformes | 0 | 1 |
| *Calidris* | Unk. sandpiper | Scolopacidae | Charadriiformes | 0 | 1 |
| *Cepphus grylle* | Black Guillemot | Alcidae | Charadriiformes | 0 | 1 |
| *Charadrius vociferus* | Killdeer | Charadriidae | Charadriiformes | 0 | 1 |
| *Gavia* | Unk. loon | Gaviidae | Gaviiformes | 0 | 1 |
| Laridae | Unk. Laridae | Laridae | Charadriiformes | 0 | 1 |
| *Larus* | Unk. gull | Laridae | Charadriiformes | 0 | 1 |
| *Larus argentatus smithsonianus* | Herring Gull | Laridae | Charadriiformes | 0 | 1 |
| *Limnodromus griseus* | Short-billed Dowitcher | Scolopacidae | Charadriiformes | 0 | 1 |
| *Melanitta deglandi* | White-winged Scoter | Anatidae | Anseriformes | 0 | 1 |
| Procellariidae | Unk. shearwater | Procellariidae | Procellariiformes | 0 | 1 |

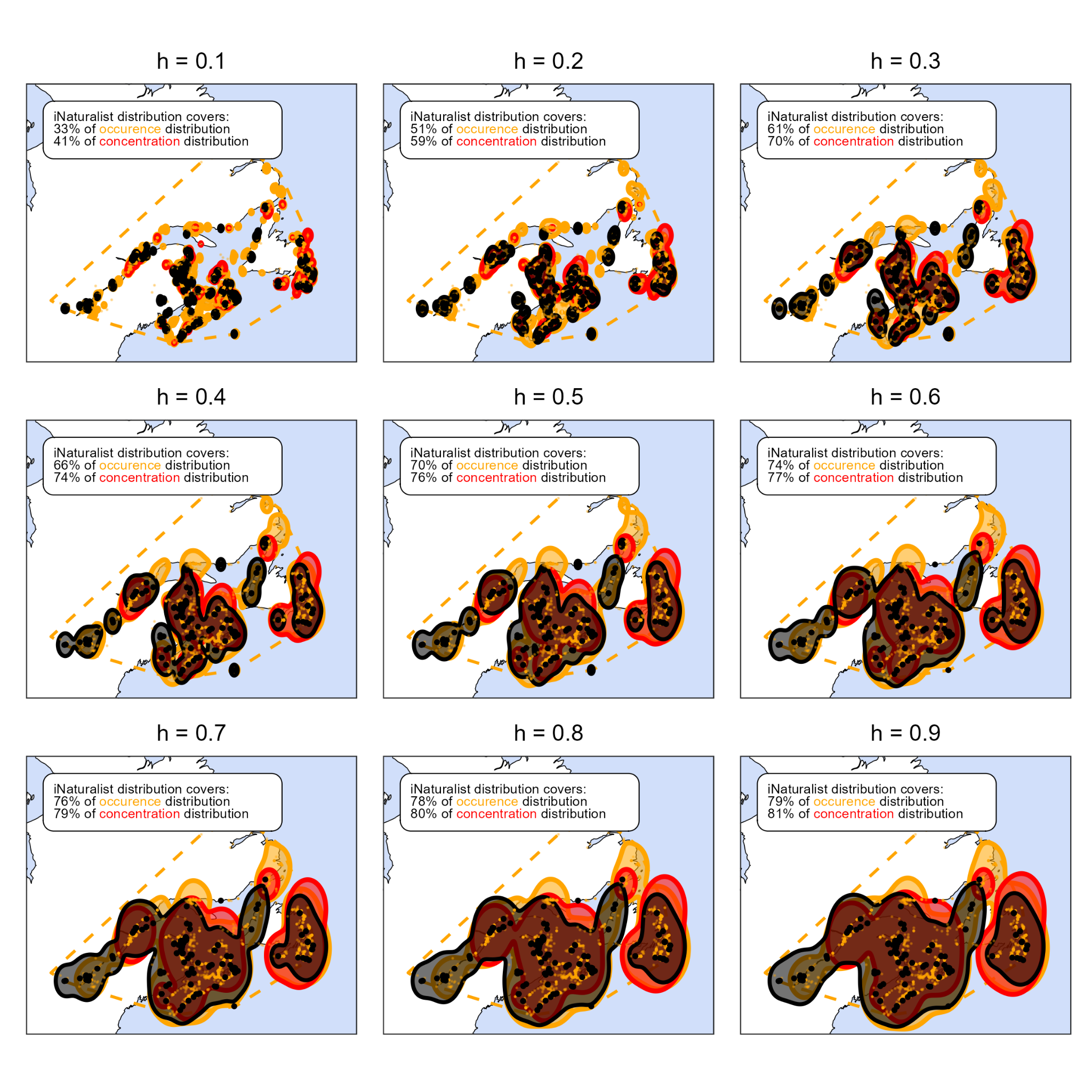

**Figure S1.** Spatial comparisons of mortality across a range of smoothing parameters (*h*) used to estimate spatial distribution (95% isopleth of mortality locations). Black contours show spatial distribution from the iNaturalist dataset. Comprehensive occurrence distribution (orange) treats mortality records from the comprehensive dataset as individual points, regardless of reported mortality, whereas comprehensive concentration distribution (red) duplicates mortalities based on the number of dead birds reported at each location. Orange dashed line indicates the full extent (convex hull) of mortality reports from the comprehensive dataset.
